# Agrochemical pollution increases risk of human exposure to schistosome parasites

**DOI:** 10.1101/161901

**Authors:** Neal T. Halstead, Christopher M. Hoover, Arathi Arakala, David J. Civitello, Giulio A. De Leo, Manoj Gambhir, Steve A. Johnson, Kristin A. Loerns, Taegan A. McMahon, Karena Nguyen, Thomas R. Raffel, Justin V. Remais, Susanne H. Sokolow, Jason R. Rohr

## Abstract

Roughly 10% of the global population is at risk of schistosomiasis, a snail-borne parasitic disease that ranks among the most important water-based diseases of humans in developing countries^1–3^. Increased prevalence, infection intensity, and spread of human schistosomiasis to non-endemic areas has been consistently linked with water resource management related to agricultural expansion, such as dam construction, which has resulted in increased snail habitat^1,4–6^. However, the role of agrochemical pollution in human schistosome transmission remains unexplored, despite strong evidence of agrochemicals increasing snail-borne diseases of wildlife^7–9^ and a projected 2- to 5-fold increase in global agrochemical use by 2050^10^ that will disproportionately occur in schistosome-endemic regions. Using a field mesocosm experiment, we show that environmentally relevant concentrations of fertilizer, the common herbicide atrazine, and the common insecticide chlorpyrifos, individually and as mixtures, increase densities of schistosome-infected snails by increasing the algae snails eat (fertilizer and atrazine) and decreasing densities of snail predators (chlorpyrifos). Epidemiological models indicate that these agrochemical effects can increase transmission of schistosomiasis. Hence, the rapid agricultural changes occurring in schistosome-endemic regions^11,12^ that are driving increased agrochemical use and pollution could potentially increase the burden of schistosomiasis in these areas. Identifying agricultural practices or agrochemicals that minimize disease risk will be critical to meeting growing food demands while improving human wellbeing^13,14^.

## Main Text

The global human population is expected to reach approximately 9.7 billion people by 2050^12^. To meet the food demands necessary to support this population, agricultural production is projected to increase 60 to 70 percent globally, with fertilizer use increasing 2- to 4-fold and pesticide use 2- to 5-fold relative to levels in 2000^10,11^. Most of the increase in both human population and agrochemical use will occur in developing regions of the world where schistosomiasis is endemic^10–12^. For example, agricultural production is expected to nearly triple in sub-Saharan Africa, the region experiencing the highest population growth rates^11^.

Schistosomiasis is caused by trematodes (flatworms) of the genus *Schistosoma* whose transmission relies on freshwater snails that act as an intermediate host. Humans (and various other mammal species) act as the definitive host (the host supporting the adult life stage of the parasite) and are infected when cercariae (the free-swimming life stage of trematodes) released from snails in infested waters penetrate through the skin of the definitive host and mature into adult worms. Global control strategies generally rely on morbidity control through treatment with praziquantel that kills adult worms harbored in human hosts, but drug therapy does not prevent re-infection from future exposure to cercariae. Thus, elimination of schistosomiasis has proven difficult throughout most of its geographic extent, with over 700 million people living in or near schistosome-endemic areas (and therefore at risk of infection)^1,15^, with at least 218 million people in need of treatment for infection as of 2015^16^.

In a trematode-amphibian system that provides a wildlife analog to the schistosome-human system, herbicides and fertilizers increased trematode transmission by stimulating the growth of attached algae (periphyton), the food source for snails (a bottom-up ecological effect)^7,8^. Hence, there is good reason to postulate that agrochemicals might also have important effects on human schistosomiasis. Additionally, insecticides can be deadly to insect and crayfish predators of snails, suggesting that they might increase the number of infected snails by increasing the overall density of snails (a top-down ecological effect)^9^, but links between insecticides and wildlife or human trematode infections have not been explored. Here we test the hypothesis that fertilizer, a common herbicide (atrazine), and a common insecticide (chlorpyrifos), individually and as agrochemical mixtures, amplify production of human schistosome cercariae through bottom-up and top-down effects on snail resources and predators, and that this in turn can increase schistosome transmission to humans. We were interested in agrochemical mixtures because they are more commonly detected in nature than individual agrochemicals^9,17,18^.

We created outdoor freshwater pond communities consisting of two snail predators (crayfish: *Procambarus alleni*, water bug: *Belostoma flumineum*), three snail species (*Biomphalaria glabrata* [native to the Neotropics, introduced to Africa; an intermediate host of *Schistosoma mansoni*], *Bulinus truncatus* [native to Africa, the Middle East, and parts of southern Europe; an intermediate host of *Schistosoma haematobium*], and *Haitia cubensis* [a non-host snail species native to the Caribbean and southeastern United States]), zooplankton, and algae in 60 1200-L mesocosms filled with 800-L of water. *Biomphalaria glabrata* was chosen because laboratory-reared snails were easily available, it is found in both South America and Africa making our results relevant to two continents, and its native range overlaps extensively with that of *H. cubensis* in the Caribbean. We included *H. cubensis* as a non-schistosome host snail species to provide a potential alternative prey source for crayfish predators rather than forcing these predators to only consume schistosome-hosting snails. Agrochemical treatments were applied to the mesocosms in five replicate spatial blocks and consisted of water and solvent (0.0625 mL/L acetone) controls, and atrazine (102 μg/L), chlorpyrifos (64μg/L), and fertilizer (4400 μg/L N and 440 μg/L P) individually and in all possible combinations (see Methods for details and additional treatments). Globally, atrazine and chlorpyrifos are among the most-used herbicides and insecticides, respectively^19–22^, and were applied at their estimated environmental concentrations calculated using US EPA software (see Methods). All three agrochemicals are used commonly in schistosomiasis-endemic regions^20–22^. Mature *S. mansoni* and *S. haematobium* eggs collected from infected Siberian hamsters were added to each mesocosm at three time points to simulate egg introduction from humans in an endemic setting. From each mesocosm, we quantified algal and snail abundance, light levels, and snail reproduction every other week; *S. mansoni* cercariae shedding rates from *Bi. glabrata* in weeks 8-10; and snail and predator densities as well as snail infection status (for *Bi. glabrata* and *Bu. truncatus*) at the end of the experiment. Additionally, we conducted toxicity tests to evaluate whether there were any ecologically relevant direct effects of the agrochemicals on the egg, miracidium, or cercaria stages of both schistosomes.

A combined factor and path analysis revealed that both top-down and bottom-up effects of the agrochemicals indirectly contributed to increases in infected *Bi. glabrata* densities through increases in overall (infected and uninfected) densities of *Bi. glabrata* (Figs. 1A, 2; Extended Data Tables 1, 2). While *Bi. glabrata* was the only snail species for which a sufficient number of infected individuals were alive at the conclusion of the experiment for analysis, treatment effects on the reproductive output and final densities of all three snail species were significant and in the same direction (Fig. 1A; Extended Data Table 1). Chlorpyrifos reduced densities of crayfish and water bugs (Fig. 1B), which indirectly increased densities of all three snail species by releasing them from predation (Fig. 1C). Both fertilizer and atrazine increased densities of all snail species (Fig. 1D) by increasing algal productivity (Fig. 1E). Fertilizer increased the densities of both suspended and attached algae (see Supplement). Consistent with previous studies^7^, atrazine decreased suspended algae and increased the photosynthetic efficiency of attached algae because the reduction in suspended algae increased light availability for the periphyton (slope between phytoplankton chlorophyll *a* and water column light: coef ± se = −334 ± 166; P = 0.0451). This indirect positive effect of atrazine on attached algae was even greater in the presence of fertilizer (Fig. 1A, Extended Data Table 1).

**Fig. 1.**
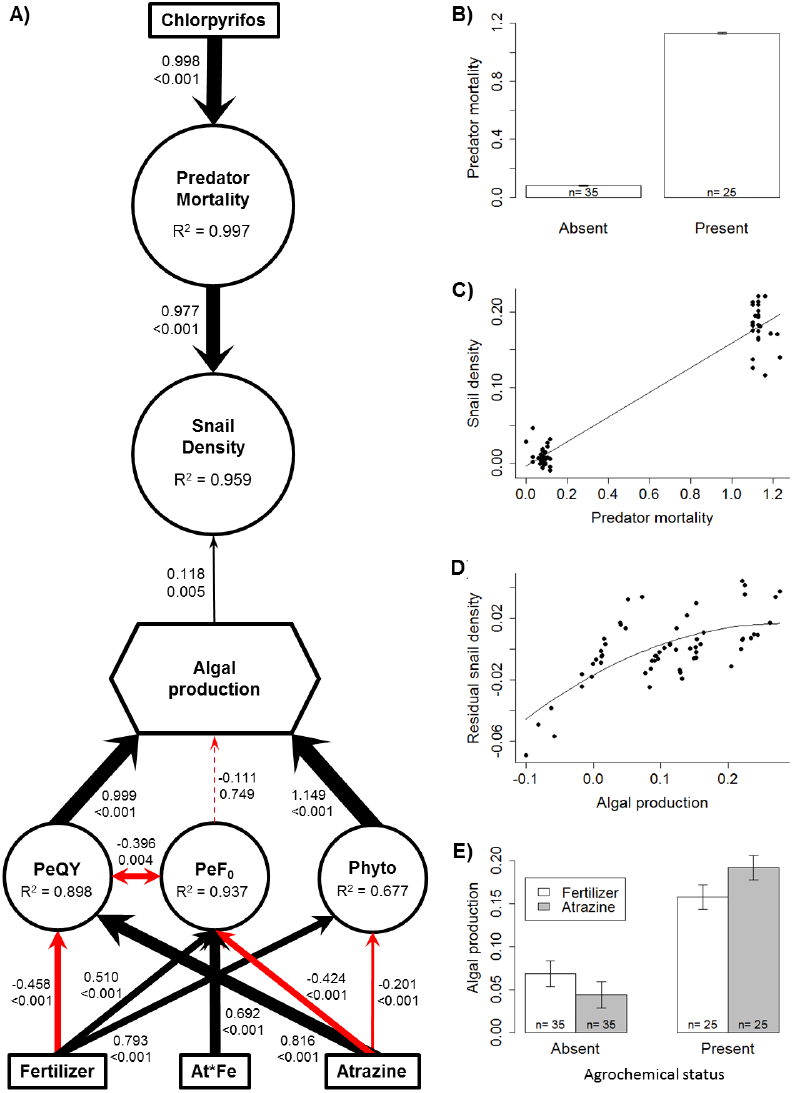
Combined factor and path analysis (**A**) demonstrating top-down effects of chlorpyrifos increasing predator mortality (**B**) and snail density (**C**) and bottom-up effects of atrazine and fertilizer increasing snail density (**D**) through increased algal productivity (**E**). Size of arrows in **A** are scaled to the standardized coefficient (top number next to each arrow), with black and red arrows indicating positive and negative coefficients, respectively. Double-ended arrows exhibit significant covariation accounted for in the structural equation model. P-values for paths in the model are reported below each standardized coefficient. Boxes represent exogenous predictor variables, circles represent latent variables, and algal production was measured as a composite variable (hexagon). Indicator variables for latent and composite variables have been omitted from the figure to reduce visual complexity, but are reported in Extended Data Table 1. Importantly, the latent variable snail density represents the densities of all three snail species at multiple life stages (egg, hatchling, and adult), all of which exhibited similar responses across treatments. Panel **E** represents the net main effects of fertilizer and atrazine presence on composite algal productivity. Axes on panels **B-E** are derived from latent variable scores for each replicate and thus have no units of measurement; however, raw data are available in the supplemental materials. PeQY = periphyton photosynthetic efficiency; PeF_0_ = periphyton chlorphyll *a*; Phyto = phytoplankton chlorophyll *a* and photosynthetic efficiency; At*Fe = atrazine ×fertilizer interaction term.

**Figure 2.**
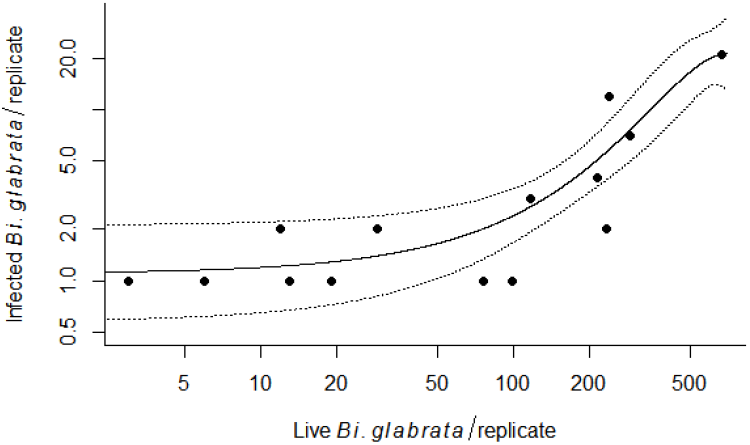
Actual number of infected *Biomphalaria glabrata* as a function of live *Bi. glabrata* at the end of the experiment. The response shown is restricted to mesocosm tanks in which infected *Bi. glabrata* were present, effectively depicting the count portion of the zero-inflated model used to analyze effects on infected *Bi. glabrata* density (see Methods and Extended Data Table 2 for full model results). Live *Bi. glabrata* density was the only significant predictor of the count of infected *Bi. glabrata* in the model (other than spatial block) and explained 89% of the variation in the density of *S. mansoni*-infected *Bi. glabrata* in replicates in which infected snails were present. The solid line represents the predictions from a generalized linear model with a Poisson distribution including first- and second-degree polynomial terms for live *Bi. glabrata* density as predictors. Dashed lines indicate the 95% confidence band.

Top-down regulation of snails by macroarthropod predators, particularly crayfish, was much stronger than bottom-up effects mediated by algal resources in this experiment (Fig. 1). This is consistent with several previous studies that showed that decapod crustaceans are effective biocontrol agents for reducing populations of *Biomphalaria* spp. and *Bulinus* spp.^4,23–27^. Separate field trials using the non-native crayfish *Procambarus clarkii* in Kenya and reintroduction of the West-African native prawn *Macrobrachium vollenhovenii* in Senegal were both successful in reducing *Bulinus* spp. densities and *S. haematobium* infection rates in humans^5,28^. Additionally, reductions in the density of a molluscivorous fish have been linked to increased infection rates and transmission of urinary schistosomiasis in humans^29,30^, underscoring the importance of predators in mediating infection dynamics in natural systems^31^.

The combined effects of agrochemicals in our mesocosm experiment accounted for 95.9% of the variation in overall snail densities accounting for all three snail species) in our path model (Fig. 1A; Extended Data Table 1). Overall final densities of *Bi. glabrata* accounted for 89.0% of the variation in the densities of infected *Bi. glabrata* (in replicates with infected snails; Fig. 2) and *Bi. glabrata* density was the only significant predictor of densities of infected *Bi. glabrata* (coef ± se = 0.0056 ± 0.0012; P < 0.001; Extended Data Table 2). Together, the indirect effects of agrochemical exposures on snail densities mediated through trophic interactions (top-down and bottom-up effects) and the effect of *Bi. glabrata* density on the density of infected *Bi. glabrata* accounted for 85.4% of the variation in densities of infected *Bi. glabrata*. Importantly, there was no evidence of direct effects of agrochemicals on the number of infected *Bi. glabrata* after controlling for *Bi. glabrata* density (Extended Data Table 2), infection prevalence (Extended Data Table 3), cercaria production per snail (Extended Data Table 4), cercarial survival (up to 12 h of exposure; Extended Data Table 5), or schistosome egg viability in toxicity trials (Extended Data Table 6).

To examine the significance of the mesocosm results for human infection, we expanded on classic^32,33^ and recent^5^ mathematical modeling studies of schistosomiasis transmission by incorporating into models the observed agrochemical effects from our mesocosm experiment, effects from previously published studies examining the same agrochemicals and endpoints as our mesocosm experiment, and parameters fit to previous research on *S. haematobium* transmission to humans in Senegal (see Methods). Our epidemiological model revealed that, in the absence of agrochemical effects and snail predators, the basic reproduction number, *R*^*0*^. (the expected number of mated female worms produced by a single mated female worm in a disease-free setting), was 3.60 (95% CI: 1.32 – 6.06; Fig. 3A), consistent with previous estimates and the endemic nature of human schistosomiasis in Senegal^5^. The addition of snail predators reduced *R0* below 1 (Fig. 3A, B), the minimum threshold for sustained transmission of the disease in the human population, supporting the notion that snail predators can reduce schistosomiasis and protect human health^5^. In contrast, by reducing snail predators, ecologically relevant concentrations of chlorpyrifos increased *R0* up to 10-fold relative to controls (Fig. 3A, B, D), suggesting that the removal of snail predators caused by pesticides may lead to a remarkable increase in disease transmission. In the absence of predators or the presence of chlorpyrifos, atrazine and fertilizer further increased *R0* and human risk by approximately 28% through bottom-up effects (Fig. 3A, C, D).

**Fig. 3.**
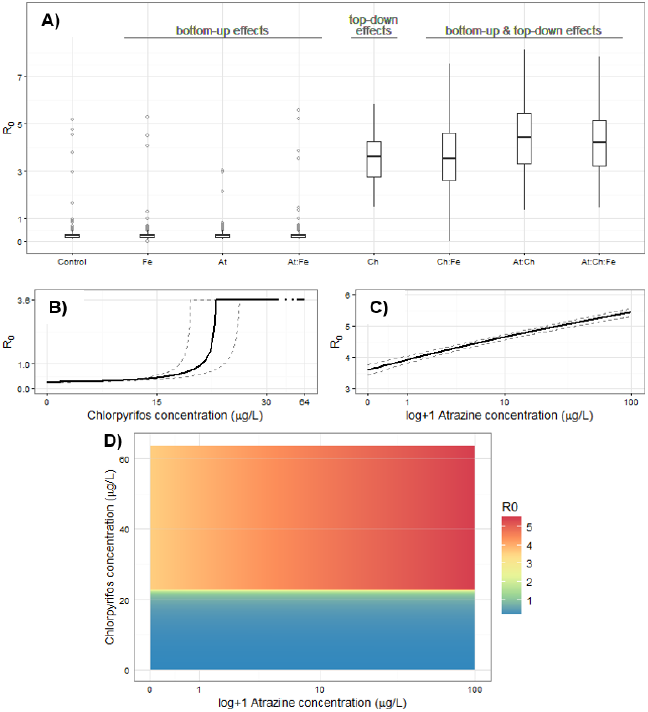
Results of a mathematical model examining the influence of each agrochemical treatment from the mesocosm experiment on estimates of *R*_*0*_ **(A)**, and drawing from other experiments that examine the same agrochemicals and affected pathways to investigate the concentration-dependent *R*_*0*_ of chlorpyrifos **(B)** and atrazine **(C)** as well as their combined influence **(D)**. Estimates of *R*_*0*_ were derived from Monte Carlo simulations that incorporate uncertainty associated with both model fitting and agrochemical parameters (see supplementary text). Boxes in **A** represent the median and interquartile range (IQR), whiskers represent values within 1.5*IQR, and outliers (points outside of 1.5*IQR) are plotted as light gray points. When chlorpyrifos is absent, transmission is restricted by top-down regulation of the snail population, causing median *R*_*0*_<1 in the control and first three treatment groups. In the presence of chlorpyrifos, median *R*_*0*_ estimates are >1, suggesting endemic transmission, and bottom-up effects from atrazine and/or fertilizer act in conjunction with top-down effects to further increase median *R*_*0*_ estimates. Maximum *R*_*0*_ in **B** is achieved when chlorpyrifos concentration is sufficient to eliminate the predator population, as in the mesocosm (64 μg/L). In a predator-free setting, *R*_*0*_ is equivalent to that estimated from the fitted model at baseline, but rises as atrazine concentration (log+1 transformed) increases due to bottom-up stimulation of the snail population. Dashed lines in **B** & **C** represent, respectively, the 95% confidence interval of predator mortality across the indicated range of chlorpyrifos concentration as estimated in^37^ and the 95% confidence interval of snail density dependence increases across the indicated range of atrazine concentrations as estimated from^38^. Expected values of *R0* driven by mixtures of atrazine acting on snail population dynamics as in^38^, and the chlorpyrifos acting on snail predators as in^37^ show that agrochemical mixtures have a pronounced influence on transmission intensity **D**.

To our knowledge, this work represents the first experimental research: 1) to examine the top-down effects of insecticides on trematode transmission; 2) to quantify the top-down and bottom-up effects of agrochemicals on the transmission of human schistosomes; 3) to establish an experimental study system on human schistosomes in outdoor mesocosms; and 4) to link experimental findings on agrochemical effects to human schistosomiasis risk by using parameterized epidemiological transmission models. Given that agrochemical use is expected to rise 2-to 5-fold globally in the next 35 years to meet growing food demands^10^, our study has important public health implications in schistosomiasis-endemic regions, as for the first time to our knowledge, it provides evidence of the potential impact of agrochemicals on the transmission of human schistosomes. These documented effects might be even more pronounced given that increases in agrochemical use in schistosome-endemic regions will likely exceed the expected 2- to 5-fold mean global increase given that human population growth rates in schistosome-endemic regions are projected to be much higher than throughout most of the more developed world^11,12^. Our results also support recent findings that the presence of generalist predators of snails, such as crayfish (tested here) and the prawn *Macrobrachium vollenhoveni* (native to western Africa) – both of which are omnivores with very similar diets^4,5,23,28,34,35^ – can limit or prevent sustained transmission of schistosomiasis (i.e. *R*_*0*_ < 1) by controlling the density of infected snails^5,24,28^. However, we also demonstrate that the common insecticide chlorpyrifos can induce considerable mortality in the snail predator population at environmentally relevant concentrations, resulting in *R*_*0*_ estimates equivalent to those in predator-free environments that experience endemic transmission (i.e. *R*_*0*_ *>* 1). Additionally, our results suggest that applications of the common herbicide atrazine and fertilizer can increase the risk of human schistosomiasis in situations where snail predators, such as prawns, exist at densities too low to effectively regulate snail populations (e.g., where dams have been constructed^4,5^). Furthermore, environmental conditions (e.g., rainfall intensity and soil characteristics) throughout most schistosome-endemic regions render surface waters highly vulnerable to pesticide runoff^36^ and projected increases in agricultural activity in these countries is likely to result in significantly higher likelihood of agrochemical contamination of surface waters. To ensure that schistosomiasis-endemic regions can address their current and pending human malnutrition crises without increasing schistosomiasis, it will be important to implement farming practices that minimize agrochemical runoff, continue advances in the sustainable use of snail biological control agents^5^, and identify agrochemicals that can increase food production without increasing snail densities.

## Methods

### Mesocosm Experiment

#### Experimental design

We established outdoor freshwater ponds in 1200-L mesocosms filled with 800 L of water at a facility approximately 20 miles southeast of Tampa, FL, USA. Tanks were filled with tap water on 4 June 2010 and allowed to age for 48 h before being seeded with algae (periphyton and phytoplankton) and zooplankton collected from local ponds on 6 June 2010. Algal and zooplankton communities were allowed to establish over a four-week period and 40 L of water was mixed between tanks weekly to attempt to homogenize initial communities before application of agrochemical treatments. Sediment (1 L play sand and 1 L organic topsoil (The Scotts Company, Marysville, OH, USA)) was added to each tank on 1 July 2010. Immediately before application of agrochemical treatments on 8 July 2010, snails (27 *Biomphalaria glabrata* (NMRI strain), 11 *Bulinus truncatus* (Egyptian strain), and 30 *Haitia cubensis*) and snail predators (3 crayfish (*Procambarus alleni*), and 7 giant water bugs (*Belostoma flumineum*)) were added to each tank. Initial snail and predator densities were chosen within ecologically relevant limits and determined by availability from either the NIAID Schistosome Resource Center (for laboratory passaged strains of *Bi. glabrata* and *Bu. truncatus*) or local availability for *H. cubensis, P. alleni*, and *B. flumineum* from wetlands in the Tampa, FL, area. The mesocosm experiment was approved under USF Institutional Biosafety Committee Study number 0971.

Sixty tanks were randomly assigned to one of 12 treatments in 5 replicated spatial blocks. Water and solvent (0.0625 mL/L acetone) controls were used to ensure that any observed effects in agrochemical treatments were not due to the presence of solvent. The herbicide atrazine and the insecticide chlorpyrifos were dissolved in acetone and applied at their respective estimated peak environmental concentrations (EEC: atrazine = 102 μg/L; chlorpyrifos = 64 μg/L), determined using USEPA’s GENEEC (v2.0, USEPA, Washington, D.C.) software, manufacturers’ label application recommendations, and the physicochemical properties of each pesticide (Extended Data Table 7). Target concentrations of fertilizer (N: 4,400 μg/L, P: 440 μg/L) were based on ponds identified as highly productive in a field survey conducted by Chase^39^. Fertilizer was applied as a mixture of sodium nitrate and sodium phosphate dissolved in acetone. Each chemical was applied individually at its EEC, at 2x the EEC, and in all pairwise combinations. The 2x EEC treatments were included as an additional reference to account for pairwise mixtures having approximately twice the amount of chemicals added. Technical-grade pesticides were used for all treatments (purity > 98%; Chemservice, West Chester, PA, USA) and actual concentrations of pesticides applied to the replicates were confirmed using ELISA test kits (Abraxis, LLC, Warminster, PA, USA) in the Rohr lab. ELISA assays were calibrated by using standards of known concentration for each pesticide, or calculated from established cross-reactivity to the chemical used to determine the standard curve. For any nominal concentrations below the limit of detection for the kit, we confirmed the concentration of the stock solution used for serial dilutions.

#### Snail infections

*Schistosoma mansoni* (NMRI strain) and *S. haematobium* (Egyptian strain) eggs were collected from infected Siberian hamsters and added to mesocosms at three separate occassions during the experiment. Eggs were added on multiple occasions after application of agrochemical treatments to better simulate the relatively constant input of schistosome eggs into waterbodies as opposed to more infrequent pulses of agrochemical runoff into surface waters. Snails and infected hamsters were provided by the NIAID Schistosomiasis Resource Center. Five infected hamsters were euthanatized on 27 July 2010, 4 August 2010, and 12 August 2010), and *S. haematobium* eggs were collected from the intestines. Eggs were isolated from tissue using a handheld immersion blender and collected on a 45 μm USA standard test sieve (Fisher Scientific, Pittsburgh, PA, USA). Mature eggs were stored in a 1.4% NaCl solution to inhibit hatching in a 50 mL centrifuge tube. Eggs were suspended repeatedly using a vortex mixer and sixty-five 3mL aliquots were prepared for each schistosome species and added to the tanks within two hours of collection. An additional three aliquots were preserved to quantify the total number of eggs added to each tank. Egg viability was quantified by placing subsamples of the remaining mature eggs in artificial spring water^40^ and observing the proportion of hatched miracidia within 1 h. The mean number of *S. mansoni* eggs in each aliquot was 981.1 (± 46.5 SEM) eggs, with a mean viability of 29.4% (± 4.6% SEM), which resulted in an estimated 289 *S. mansoni* miracidia added to each tank at each weekly addition. The mean number of *S. haematobium* eggs in each aliquot was 2,276.7 (± 107.5 SEM) eggs, with a mean viability of 8.5% (± 1.9% SEM), which resulted in an estimated 193 *S. haematobium* miracidia added to each tank at each weekly addition. Collection of schistosome eggs from infected hamsters was approved by animal care and use committee protocols T 3829 and R 3517 at the University of South Florida.

#### Biosafety precautions

Strict biosafety protocols were established and approved by USF Biosafety (IBC #1334) to minimize the risk of infection to researchers and escape of snails from the mesocosms. Researchers working at the mesocosm facility wore personal protective equipment, including shoulder-length PVC gloves (#7451, Galeton, Foxborough, MA, USA), when removing or replacing items in tanks. In addition, all researchers working on the mesocosm experiment had blood drawn before and several months after conducting the experiment. Blood samples were sent to the Centers for Disease Control and Prevention to test for schistosome infections. Tanks had an inward-projecting outer rim along the top edge, were only filled halfway, and were covered with heavily-weighted shade cloth to prevent snail escape or entry of any large organisms. The mesocosm facility was surrounded by two layers of silt fence with molluscicide (1.0% iron phosphate; Natria®, Bayer Advanced, Research Triangle Park, NC, USA) applied between the fences at the recommended rate of 1 pound per 1,000 square feet every two weeks during the experiment. Tanks were a minimum of 200 m from the nearest waterbody and the entire facility was surrounded by chain link and barbed wire fencing. At the end of the experiment, each tank was over-treated with pool shock (71.8% trichloro-s-triazinetrione, Recreational Water Products, Buford, GA, USA; applied at 0.15 g/L) to kill all of the snails and schistosomes before the tanks were emptied and the snails were removed and preserved.

#### Data collection

Periphyton measurements were recorded from 100 cm^2^ clay tiles suspended vertically 15 cm from the bottom of each tank (approximately 20 cm below the water’s surface), facing south along the northern wall of each tank. Five clay tiles were added to each tank when they were initially filled with water. Algal samples were collected immediately prior to agrochemical addition (Week 0) and at 1, 2, 4, 8, and 12 weeks post-application. Phytoplankton and periphyton chlorophyll *a* and photosynthetic efficiency (measured as F0 and QY, respectively), were measured from samples stored in darkness for 1 h, using a handheld fluorometer (Z985 Cuvette AquaPen, Qubit Systems Inc., Kingston, Ontario, Canada). Temperature and light levels were quantified on the same dates as algal sampling by suspending a data logger (HOBO Pendant UA-002-64, Onset Computer Corporation, Bourne, MA, USA) 20 cm below the water surface for 30 minutes in each replicate within a spatial block near midday. Loggers were rinsed in tap water after the 30 minute data collection period for each block before being transferred to the next spatial block to avoid cross contamination of agrochemicals. Midday temperatures 20 cm below the water surface were 32.55±0.10°C on 9 July 2010, 30.49±0.08°C on 15 July 2010, 31.47±0.10°C on 22 July 2010, and 31.29±0.07°C on 5 August 2010 (all mean ± se).

Snail reproductive effort and density was estimated using two 15 × 30 cm pieces of Plexiglass placed in each tank; one suspended vertically 10 cm from the bottom of each tank and one resting horizontally along the tank bottom. Snail egg masses, juveniles, and adults were quantified from each sampler at weeks 1, 2, 4, 8, and 12. Visual searches for dead *P. alleni* and *B. flumineum* occurred 24 and 48 h after agrochemical addition, and upon each snail sampling session. Ten weeks after agrochemical addition, pool shock was added to each tank as described above to kill any infective schistosome cercariae and tanks were subsequently drained through a kick net (800/900 μm, 425-K11, Wildlife Supply Company, Yulee, FL, USA) to collect remaining organisms. All snails and macroinvertebrates were collected, fixed in formalin for one week, and subsequently preserved in 70% ethanol. Snail infection status was determined by cracking each snail’s shell and inspecting the hepatopancreas and gonads under a dissecting microscope.

#### Data analysis

There was no effect of solvent on any response variables, so solvent and water controls were pooled and treated as a single control treatment. Likewise, there was no effect of concentration on any of the observed response variables, so 1x and 2x EEC single pesticide treatments were combined for analysis. Photosynthetic efficiency was logit-transformed prior to analysis. All other response variables were natural log+1 transformed.

Structural equation modeling was used to explore combined causal pathways of pesticide mixtures using the lavaan package^41^ in R statistical software^42^. Because a sample size of 60 tanks restricted the number of causal pathways we could infer, we first constructed a latent variable for predator mortality (*P. alleni* and *B. flumineum* mortality at 24h and the end of the experiment) and a second model consisting of latent variables for phytoplankton production (F0 and QY from weeks 1-8), periphyton chlorophyll *a* (F0 from weeks 1-4), and periphyton photosynthetic efficiency (QY from weeks 1-4). Model comparison using AICc was performed to select the best latent variable model from alternative configurations of indicator variables (i.e., algal parameters as separate or combined latent variables without a composite variable). The scores for each latent variable model were then extracted using the predict function and used for construction of a structural equation model that included snail response variables (number of egg masses on snail samplers from weeks 1-4, the number of snail hatchlings on snail samplers from weeks 4-8, and the number of live snails collected at the end of the experiment for each species, including the non-host *H. cubensis*) as indicators of a latent variable of overall snail density.

The relationship between infected *Bi. glabrata* density and density of live *Bi. glabrata* at the end of the experiment was analyzed using generalized linear models in the pscl package^43,44^ in R statistical software^42^. This relationship is only presented for *Bi. glabrata* because there were too few infected *Bu. truncatus* alive at the end of the experiment to perform the same analysis. Final *Bi. glabrata* density in addition to fixed main effects of agrochemicals, their interactions, and block were used as predictors of the count of infected *Bi. glabrata* in each tank, with a zero-inflated Poisson distribution (Extended Data Table 2). We also tested for direct effects of agrochemicals on infection prevalence using a beta binomial error distribution of infected vs uninfected *Bi. glabrata* in each tank with fixed effects of agrochemicals, their interactions, and block as predictors (Extended Data Table 3). Model selection indicated that the beta binomial error distribution was a better fit to the prevalence data than a binomial distribution (ΔAICc = 8.3). Analysis of prevalence data was performed using the glmmADMB package^45,46^ in R^42^. Light availability was tested as a response of periphyton chlorophyll *a* and fixed effects of block in week 2, using the glmmADMB package in R.

#### Effects of atrazine and fertilizer on algal dynamics and snail densities

The observed algal dynamics are consistent with previous research^7^. Fertilizer increased phytoplankton density (chlorophyll *a*) and photosynthetic efficiency, and increased periphyton density, but photosynthetic efficiency of periphyton was reduced in fertilizer treatments. However, phytoplankton density was a negative predictor of light availability in the water column (coef ±se = −334 ± 166; P = 0.0451), and decreased light availability is therefore likely to reduce the photosynthetic efficiency of periphyton. Conversely, atrazine decreased phytoplankton chlorophyll *a* and photosynthetic efficiency. Thus, although atrazine negatively impacted periphyton chlorophyll *a*, photosynthetic efficiency of periphyton increased in the presence of atrazine because more light was available for photosynthesis, and a positive interaction between the joint presence of atrazine and fertilizer in mixtures increased periphyton density substantially (Extended Data Fig. 1, Extended Data Table 1). The lack of complex refugia for snails in the mesocosms may have artificially decreased the apparent strength of bottom-up effects on snail densities relative to top-down regulation by predators. We explored the potential for a submerged macrophyte, *Hydrilla verticillata*, to provide refugia for snails in a separate mesocosm experiment, and found no evidence that *H. verticillata* provided effective refugia from omnivorous crayfish (see *Effects of Snail Refugia and Alternative Food Sources for Predators on Snail Densities* below). However, because crayfish readily consume both living and decaying plant matter, non-consumable refugia might provide a stronger mediating effect on the relative strengths of top-down versus bottom-up regulation of snail populations.

### Cercaria Production Experiment

*Aim:* To test for indirect effects of agrochemical exposure on cercarial production (through potential effects of agrochemicals on resource composition, quality, and/or abundance).

#### Experimental design

Fifteen freshwater mesocosms were established at the same time and using the same methods as noted above for the main mesocosm experiment, with the exception that no snails or snail predators were added to these tanks. Instead, algal and zooplankton dynamics were allowed to respond to agrochemical treatments in the absence of periphyton herbivores. This mesocosm experiment was also approved under USF Institutional Biosafety Committee Study number 0971.

Tanks were randomly assigned to one of 5 treatments in 3 replicated spatial blocks. Atrazine, chlorpyrifos, and fertilizer were applied at their respective EECs as described previously. Solvent controls were used to account for the presence of solvent used to deliver agrochemicals in solution. In addition, a treatment combining atrazine and fertilizer at their respective target concentrations was included to test for a potential interaction between these two agrochemicals. Technical-grade pesticides were used for all treatments and actual concentrations of chemicals applied to the replicates were confirmed as described above for the main mesocosm experiment.

Forty infected *Bi. glabrata* (NMRI strain) were obtained from the NIAID Schistosomiasis Resource Center and added to the first block of tanks four weeks post-miracidia exposure (snails exposed 28 July 2010, added to first block 24 August 2010). Eight snails were added to each of the five replicate tanks in one block and left in the tanks for three days before calculating cercaria production rates. On the third morning after adding snails to each tank, snails were removed from each tank and placed individually in 250 mL specimen containers filled with 100 mL of ASW for 1 h. After 1 h, snails were removed from each container and five drops of Lugol’s iodine were added to preserve and stain cercariae. Snails were haphazardly assigned to replicate tanks in the next spatial block and left for three days before repeating the cercaria production trials. This process continued until the infected snails were rotated through each block a total of two times. Cercariae were counted in the laboratory under a dissecting microscope.

#### Data analysis

We tested for main effects of agrochemical treatment on the total number of cercariae shed per hour with a negative binomial generalized linear model, using the glmmADMB package in R^45,46^. We used fixed main effects for each chemical and days since miracidia exposure and included random effects of tank nested in block nested by trial number (first or second trial) as predictors of the cercaria shedding rate. In addition, we tested for an interaction between atrazine and fertilizer in the absence of chlorpyrifos. Cercaria production rates per snail increased with increasing time since miracidial exposure (coef ±se = 0.138 ± 0.035; P < 0.001; Extended Data Table 4), but there were no effects of agrochemical exposure (all P > 0.35; Extended Data Table 4).

### Direct Effects of Agrochemical Exposure on Schistosome Cercariae

#### Aim

To test for direct effects of agrochemical exposure on the cercariae of *S. mansoni*.

#### Experimental Design

Six replicates of four agrochemical treatments (atrazine, chlorpyrifos, fertilizer and a solvent control each at their EEC as described above for the mesocosm experiment) were randomly assigned to the wells of a 24-well tissue culture plate (Falcon® # 353047, Corning Incorporated, Corning, NY, USA) containing freshly collected *Schistosoma mansoni* (NMRI strain) cercariae in 400 μL of COMBO^47^ (8.65±0.64 cercariae/well). One-hundred μL of stock solution of each agrochemical was added to randomly assigned wells at the beginning of each trial to reach the target EEC for each treatment at a total volume of 500 μL. Survival of cercariae was assessed at 2, 4, 8, 12, and 24 h after agrochemical addition using a separate 24-well plate for each time point. At the given end point for each trial, the number of dead cercariae was determined by adding 15 μL of trypan blue, a selective stain that is taken up only by dead cercariae^48^, to each well. Following staining with trypan blue, 20 μL Lugol’s iodine was added to each well to kill and stain all cercariae in the well.

#### Data Analysis

Cercarial survival at each time point was tested using a binomial generalized linear model in the glmmADMB package in R^45,46^. Fixed main effects of each agrochemical and time were used as predictors of the proportion of dead cercariae in each well. There was a significant negative effect of time since agrochemical exposure on cercarial survival (coef ± se = −0.264 ± 0.017; P < 0.001), but not of any agrochemicals (all P ≥ 0.10). When analyzing cercarial survival at each time point independently, no main effects of agrochemicals were evident within 12 h of exposure to agrochemicals (all P ≥ 0.10; Extended Data Table 5). Chlorpyrifos and fertilizer each had significant negative effects on cercarial survival at 24 h post-exposure (Extended Data Table 5). However, because infectivity of *S. mansoni* cercariae declines rapidly and is very low beyond 8-15 h^49^, any treatment effects of agrochemicals after 12 h are less ecologically relevant than earlier time points.

### Schistosome Egg Viability Experiment

#### Aim

To test for direct effects of agrochemical exposure on the egg viability of *S. mansoni* and *S. haematobium*.

Eggs were collected from the tissues of two *S. mansoni*-infected Swiss-Webster mice and two *S. haematobium-*infected Siberian hamsters, on 1 Sep 2011, 6 September 2011, and 8 September 2011. See Supplementary Methods for the mesocosm experiment for detailed methods on egg collection. Eggs were stored in 1.4% NaCl to prevent hatching before beginning egg viability trials on each day. For each species on each date, twelve agrochemical treatments (described above) were randomly applied to wells filled with 1.0 mL ASW in two spatial blocks in a 24-well tissue culture plate (Falcon® # 353047, Corning Incorporated, Corning, NY, USA). After applying agrochemicals to each well, approximately 20 eggs of either *S. mansoni* or *S. haematobium* were added to each well. The number of miracidia in each well was counted after 1 h. Lugol’s iodine was then added to each well to stain and count the unhatched eggs in each well. One plate trial was performed on each date for each species, for a total of six replicate trials per species. Egg viability was tested with a beta binomial generalized linear mixed-effects model, using the glmmADMB package in R^45,46^. Fixed main effects of and interactions between agrochemicals and random effects of block nested within plate were used as predictors of hatching success. No main effects of agrochemicals or interactions between agrochemicals on schistosome egg viability were evident (all P > 0.05; Extended Data Table 6).

### Effects of Alternative Food Sources for Predators and Snail Refugia on Snail Densities

*Aim:* To explore the potential for alternative food resources for omnivorous predators and the presence of refugia on snail densities.

We used data from a follow-up mesocosm experiment on the individual effects of different classes of herbicides and insecticides to opportunistically explore the potential for a submerged macrophyte, *Hydrilla verticillata*, to influence snail densities by providing potential refugia for snails, and providing an alternative resource for the omnivorous crayfish predator, *Procambarus alleni*.

#### Experimental design

We established 70 outdoor freshwater ponds in 800-L mesocosms filled with 500 L of water at a facility approximately 20 miles southeast of Tampa, FL, USA. Tanks were set up as previously described for the mesocosm experiment, with the addition of 5 rooted shoots of *Hydrilla verticillata* added to the sediment of each tank on 5 July 2011. Immediately before application of agrochemical treatments on 11 July 2011 (Week 0), snails (21 *Bi*. *glabrata* (NMRI strain), and 12 *Bu. truncatus* (Egyptian strain), provided by NIAID Schistosomiasis Resource Center) and snail predators (2 juvenile crayfish (*Procambarus alleni*), 8 giant water bugs (7 *Belostoma flumineum* and 3 *Lethocerus* sp.) collected from local ponds) were added to each tank. The mesocosm experiment was approved under USF Institutional Biosafety Committee Study number 0971, with the same biosafety precautions as described above.

Tanks were randomly assigned to one of fourteen treatments (six herbicides at their respective EEC, six insecticides at their respective EEC, solvent control (0.0625 mL/L acetone), and water control) in five replicated spatial blocks. All pesticides were dissolved in acetone and applied at their respective estimated peak environmental concentrations as described above.

#### Data Collection

Data were collected at intervals as described above for the previous mesocosm experiment until the conclusion of the experiment after 12 weeks on 31 September 2011).

#### Data Analysis

Because herbicides either eliminated or reduced growth of *H. verticillata* but had no apparent direct effects on invertebrate predators and insecticides eliminated invertebrate predators but had no direct effects on *H. verticillata*, we opportunistically explored the effects of *H. verticillata* and predator presence or absence on snail densities at the end of the experiment. Fixed main effects of *H. verticillata* biomass at the end of the experiment, *P. alleni* presence/absence at the end of the experiment, spatial block, and all interactions were used as predictors of the count of adult *Bi. glabrata* or *Bu. truncatus* in each tank at the end of the experiment, using a generalized linear model with a Poisson error distribution in R^42^.

#### Results

There were significant main effects and a significant interaction between *H. verticillata* biomass and crayfish presence on the densities of both snail species (Extended Data Fig 2, Extended Data Table 8). For both *Bi. glabrata* and *Bu. truncatus*, crayfish presence had a strong negative effect on adult snail densities at the end of the experiment. The densities of both snail species increased with increasing *H. verticillata* biomass in the absence of crayfish, but this effect disappeared or reversed in the presence of crayfish (Extended Data Fig 2). These results suggest that *H. verticillata* serves as an additional substrate for epiphytic algae which snails consume, and the macrophyte did not alter crayfish predation rates on snails (Extended Data Fig 2). In fact, there was no recruitment to the adult snail population when crayfish were present regardless of the density of the macrophyte.

#### Effects of prey-switching by generalist predators

Both crayfish (*Procambarus* spp.) and prawns (*Macrobrachium* spp.) are omnivorous species that consume large amounts of living and decaying plant and algal matter in addition to a wide variety of animal prey^4,5,23,25,28,35,50,51^. Therefore, the presence of alternative food resources could impact the strength of the top-down effects on snail densities^51^. We attempted to avoid forcing crayfish to consume only *Schistosoma*-harboring snail hosts in the first mesocosm experiment by adding a third, non-host snail species. In this separate mesocosm experiment, *Hydrilla verticillata* served as an alternative food resource for crayfish, yet it did not limit the effects of predation on snail densities (Extended Data Figure 2), suggesting that at least this macrophyte does not reduce the strength of top-down effects of crayfish on these snail populations and that our results are robust to the presence of alternative food resources for omnivorous snail predators. Additionally, our mathematical model (see *Modelling Experiments* below) implicitly accounts for prey-switching behavior of generalist snail predators by the use of a Holling type III functional response, which specifies that the per-capita, per-predator predation mortality of the snails is not constant but rather increases with snail density to represent how a predator’s preference switches away from snails when snail abundance is low. Finally, field research in Senegal involving the reintroduction of *M. vollenhovei* to natural waterbodies, which presumably included a much wider variety of available food resources for prawns, found very similar effects on the densities of *Bulinus truncatus* and *Bu. globosus*, reinforcing our conclusion that the top-down effects of snail predators are very strong in natural settings when predators are present.

#### Effects of refugia

The potential effects of refugia on final snail densities (and infected snail densities) is less clear from our results. Although submerged macrophytes might provide snails with a potential refuge from predators^52^, there was no evidence from the mesocosm experiment that the density of *H. verticillata* affected snail consumption rates by crayfish. However, the effects of refugia on snail population dynamics in natural environments are complex because refuge-seeking behaviors, and growth and reproduction rates of snails can be modified by the types and densities of predators, snail densities, resource availability, and snail infection status^27,53–56^. While a greater availability of refugia in the mesocosm experiments would likely have resulted in higher overall snail densities, the net effects of refugia on infected snail densities are more difficult to predict because prawns (and perhaps crayfish) show a preference for consuming schistosome-infected *Bi. glabrata* and *Bu. truncatus* in predation trials, and infected *Bi. glabrata* and *Bu. truncatus* exhibit less frequent and slower movement and lower refuge use (above water or under substrate) than uninfected snails in response to predation cues in laboratory trials^27^. The effects and interactions of refuge availability, resource availability (especially as modified by agrochemical contamination), and multiple types of predators on schistosome-infected snail densities clearly deserves further examination. However, while these factors are might affect the relative strengths of bottom-up vs. top-down regulation of infected snail densities, multiple field observations and manipulations in natural settings^4,5,25,28–31^ are consistent with our experimental and modelling results that highlight the importance of predators in regulating snail densities.

### Modelling Experiments

#### Mathematical model

A model expanding on previous^5^ and classic^32,33^ work was used to investigate agrochemical effects—of atrazine, chlorpyrifos, and fertilizer—on human schistosomiasis transmission intensity. The model includes snail population dynamics of *Bulinus truncatus*, the intermediate host of *Schistosoma haemotobium*, subject to logistic population growth and the influence of predation. We focus on *S. haematobium* for our modelling because it is the predominant schistosome species found in the village in Senegal where the epidemiological data used to parameterize our model were collected. The population dynamics of generalist predators (*P*) are included, subject to an agrochemical-sensitive mortality rate, *μ*_*p,q*_, that reflects chlorpyrifos toxicity to the predator population as estimated by the mesocosm experiments and previous work^37^. Because crayfish and prawns are generalist predators and will switch to other resources when snail densities are low, the model assumes that predator population dynamics are independent of snail density. Also included is a parameter representing agrochemical enhancement of the snail carrying capacity, *φ*_*N,q*_, which models the snail population response to bottom-up effects caused by algal stimulation by atrazine and fertilizer as estimated in the mesocosm experiments and other experiments examining the same agrochemicals and outcomes. Additional model state variables represent susceptible, exposed and infected snails (*S, E*, and *I*, respectively) and the mean worm burden in the human population (*W*). The number of mated female worms, *M*, is estimated assuming a 1:1 sex ratio and mating function, *γ*(w,k), as in ^57^. The per capita snail predation rate by predators, modelled as a Holling type III functional response as in ^58^, *Ψ* and the total snail population, *N*, are shown separately for clarity. Parameter values, definitions and reference literature are listed in Extended Data Table 9.

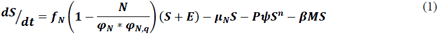

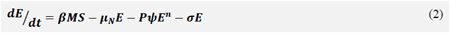

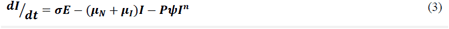

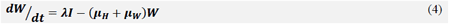

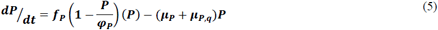

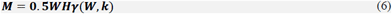

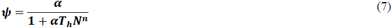

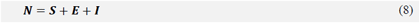

#### Derivation of R_0_

Using the next generation matrix method^59^, we calculate *R_0_* for this system as:

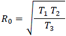

Where:

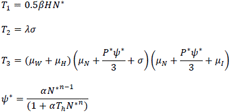

and *N*^*\**^ and *P*^*\**^ represent disease-free equilibrium values of the snail and predator populations, respectively, derived by setting equations (1) and (5) equal to 0 such that:

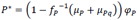

and *N*^*\**^ is estimated by solving for *N\** in the polynomial:

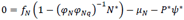

### Model fit to epidemiological data

The model expressed as equations (1) through (8) was fit to previously published worm burden data from a baseline and follow-up survey of *Schistosoma haemotobium* infection in a rural community upstream of the Diama Dam in Senegal^5^. The human-to-snail transmission parameter, *β*, and two values of the snail-to-human transmission parameter, *λ*_*lo*_ and *λ*_*hi*_, were fit to the seasonal reinfection data using maximum likelihood estimation in R with the *optim* function^42^, with all other parameters held to values shown in Extended Data Table 9, and agrochemical and predation effects turned off (i.e. *φ*_*N,q*_ = 1 and P= 0). Estimates of uncertainty associated with model fitting were generated by exploring the three-dimensional parameter space around the best-fit values of *β*, λ_*lo*_, and λ_*hi*_ (shown in Extended Data Table 9) by varying each plus or minus 90% of its best-fit value. Assuming the negative log likelihood profile follows a chi-square distribution with three degrees of freedom, all parameter triplets that have negative log likelihood within 7.815 (95% CI, two-sided chi-square critical value) of the negative log likelihood produced by the best-fit values are within the 95% confidence interval. These parameter triplets were used in Monte Carlo simulations described further below to generate estimates of *R*_*0*_. When estimating steady-state transmission indices such as *R*_*0*_ a time-weighted average of λ_*lo*_ and λ_*hi*_ was used.

### Estimating agrochemical response parameters

#### Top-down effects

Point estimates of the baseline daily predator mortality rate, *μ*_*p*_, and the chlorpyrifos-enhanced predator mortality rate, *μ*_*p,q*_, were derived directly from 24 h mortality endpoints in the mesocosm experiment by treating all *Procambarus alleni* in chlorpyrifos tanks (75 total) as a treatment cohort and all *Procambarus alleni* in chlorpyrifos-free tanks (105 total) as a control group (Extended Data Table 9). A parametric distribution of the predator mortality rate was obtained by fitting beta distributions to 5000 bootstrapped samples of daily predator mortality in each of the 25 mesocosm tanks with and 35 mesocosm tanks without chlorpyrifos added (Extended Data Table 10).

To investigate the influence of a broader range of chlorpyrifos concentrations on estimates of *R*_*0*_, a probit model of predator mortality was derived spanning the range of chlorpyrifos concentrations (0 – 64 μg/L) tested in ^37^, conservatively assuming 99% mortality in the highest tested concentration groups instead of 100% to account for potential resistance in a small number of predators. Daily, per capita predator mortality rates were derived across the range of tested concentrations and used in the *R*_*0*_ expression to generate Fig 3B-D.

Because crayfish are generalists, we modeled their predation of snails using a Holling type III functional response (eqn 7) in which the per capita predation rate is sigmoidal due to prey switching at low snail densities and restriction by the handling time (*T*_*h*_) at high snail densities^58,60,61^. Though not directly interpretable, the exponent, *n*, of the type III functional response is often assumed to be 2 for invertebrate predators^60,62^. However we also tested a range of values from 1-4 for the exponent, *n*, and found little qualitative difference in results when *n* > 1. When *n* = 1, the functional response reduces to a Holling type II in which the predation rate increases rapidly at low prey density and asymptotes at high prey densities where predation is restricted by the handling time. In our model, this leads to predation-induced extirpation of the snail population and *R*_*0*_ = 0, a result we would not expect in real-world transmission settings in which we expect prey switching by the predator population as well as refuge-seeking by snails to diminish predatory activity at decreasing snail densities.

#### Bottom-up effects

Bottom-up effects in the mesocosm were introduced through model parameter *φ*_*N,q*_, a scalar that represents a proportional change in the baseline snail density-dependence parameter, *φ_N_*. To quantify the effect of atrazine and fertilizer, alone and in combination, on this parameter while controlling for the strong influence of predators on snail population dynamics, we calculated the mean proportional increase in final *Bu. truncatus* density in the mesocosm tanks when fertilizer and atrazine were added in combination with chlorpyrifos (Extended Data Table 10). We found no previous studies in which chlorpyrifos had a significant direct effect on snail population dynamics nor on algal dynamics at the concentrations tested in the mesocosm. The distribution of the density dependence scalar, *φ*_*N,q*_, was obtained by fitting normal distributions to 5000 bootstrapped samples of the parameter estimate derived from individual tanks within each treatment group (Extended Data Table 10). To further investigate the influence of atrazine at concentrations below the maximum expected environmental concentration—as was tested in the mesocosm—we derived another atrazine-dependent scalar of the carrying capacity based on the results of ^63^. Briefly: the scalar was calculated according to the proportional increase in the peak growth rate at the tested atrazine concentrations over the observed peak growth rate of the control group, as discussed in ^38^.

#### Monte Carlo simulation

To produce estimates of *R*_*0*_ that incorporate uncertainty associated with both model fitting and agrochemical parameterization, we ran 1,000 Monte Carlo simulations for each agrochemical treatment and the control group; drawing randomly from the agrochemical parameter probability distributions described above and from the range of best-fit transmission parameters (Extended Data Table 10). The probability of sampling particular transmission parameter triplets was weighted by a normalized index of their likelihood so that triplets that better fit the model were more likely to be included in the simulation.

---

**Supplementary Information** is linked to the online version of the paper at www.nature.com/nature.

## Acknowledgments

We thank G. Agemy, Z. Babwani, J. Cook, E. Cooper, A. Earls, A. Gilbert, J. Jones, M. Kepner, S. Kilgore, B. Mathew, M. McGarrity, J. Rivera, A. Rodríguez, A. Tapilyai, and C. Towne for assistance with collecting data in the field and laboratory. Snails and infected hamsters were provided by the NIAID Schistosomiasis Resource Center. This research was supported by grants from the National Science Foundation (EF-1241889), National Institutes of Health (R01GM109499, R01TW010286), US Department of Agriculture (NRI 2006-01370, 2009-35102-0543), and US Environmental Protection Agency (CAREER 83518801) to J.R.R., and National Institutes of Health (K01AI091864) and the National Science Foundation (EAR-1646708, EAR-1360330) to J.V.R.; a University of Florida Research Innovation Award to J.R.R. and S.A.J.; and an Oakland University Research Excellence Fund award to T.R.R. S.H.S. and G.D.L have been supported by NSF CNH grant # 1414102, the Bill and Melinda Gates Foundation, NIH Grant 1R01TW010286-01 and Stanford GDP SEED grant 1183573-100-GDPAO.

## Author Contributions

J.R.R. conceived the experiment. N.T.H. and J.R.R. designed the experiment. S.A.J. provided field and laboratory facilities. N.T.H., T.A.M., K.P., K.N., T.R.R., and J.R.R. conducted the experiment. K.P. determined infection status of snails. N.T.H. and D.J.C. conducted the statistical analyses. C.M.H., A.A., M.G. and J.V.R. conducted the mathematical modeling and risk analysis. G.D.L. and S.H.S. participated to the calibration of the model and to the analysis of model results. N.T.H., C.M.H. and J.R.R. wrote the manuscript and all authors contributed to its editing.

## Author Information

Reprints and permissions information is available at <u>www.nature.com/reprints.</u> Correspondence and requests for materials should be addressed to N.T.H.

## Extended Data

Extended Data Figures 1-2

Extended Data Tables 1-10

**Extended Data Figure 1.**
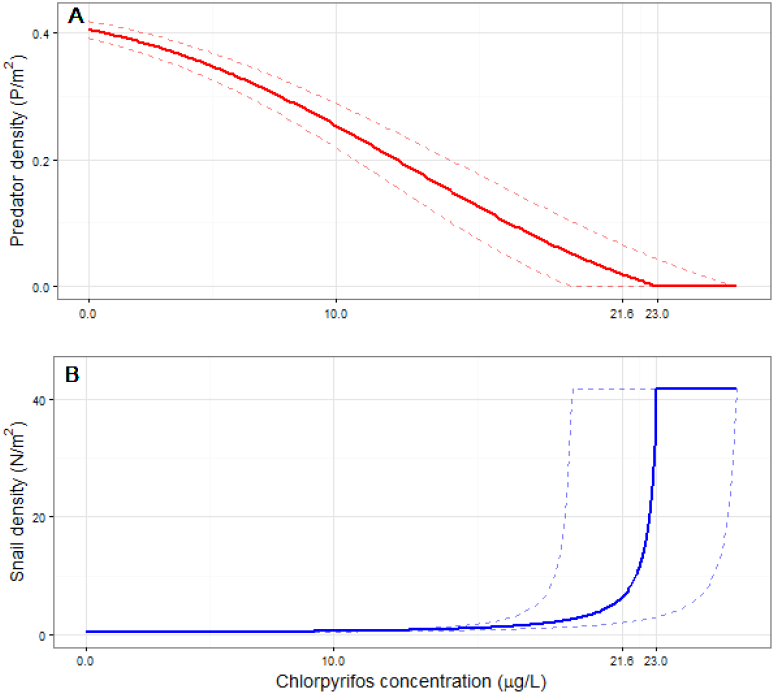
Equilibrium predator (**A**) and snail (**B**) densities at chlorpyrifos concentrations <25 ¼g/L. Interactions between these populations determine *R*_*0*_ at concentrations <23.0 ¼g/L whereas predator elimination—and endemic transmission equivalent to that in a predator-free setting—is expected to occur at concentrations >23.0 ¼g/L. At chlorpyrifos concentrations >21.6 ¼g/L, predator mortality is high enough to allow sufficient snail reproduction to occur, causing R0 >1.

**Extended Data Figure 2.**
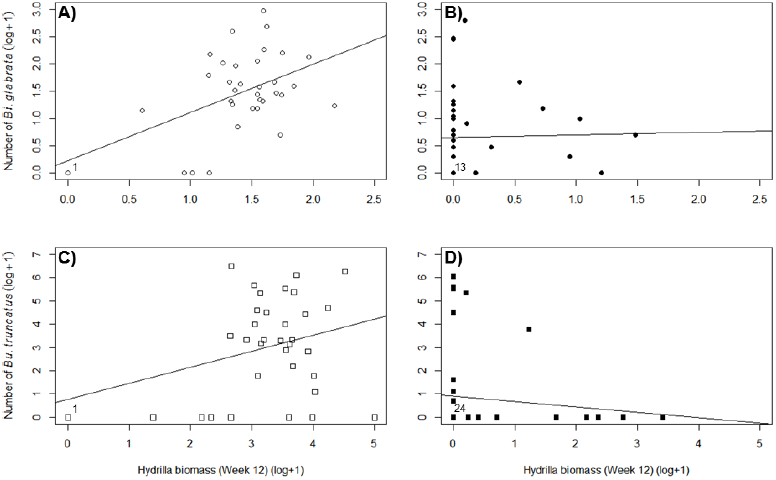
Final densities of *Biomphalaria glabrata* (circles; **A,B**) and *Bulinus truncatus* (squares; **C,D**) in response to the biomass of *Hydrilla verticillata* and predator absence (open symbols; **A,C**) or presence (solid symbols; **B,D**) at the end of a separate mesocosm experiment that included submerged macrophytes as a refugia and food source for snails and an alternative food source for omnivorous crayfish *Procambarus alleni*. For both species of snails, density increased with increasing biomass of *Hydrilla* when crayfish were absent from mesocosms, but not in the presence of predators. Thus, the density of this macrophyte species did not significantly affect the interaction strength between this crayfish predator and either *Bi. glabrata* or *Bu. truncatus*. Numbers next to 0,0 points indicate the number of replicates with those values.

**Extended Data Table 1.** Results of the structural equation model conducted in the package lavaan in R revealing the relationships among agrochemical mixtures, predator density, algal productivity, and snail density. Densities of all three snail species, including measures of reproductive effort (eggs and hatchlings) contributed significantly to the latent variable “snail density” and generally covaried positively, emphasizing the similarity of treatment effects across all snail species.

| Latent variables | Estimate | Std.err | Standardized estimate | Z-value | P (>\|z\|) |
| --- | --- | --- | --- | --- | --- |
| Predator mortality |  |  |  |  |  |
| *P. alleni* (24h) | 1.000 |  | 0.874 |  |  |
| *P. alleni* (end) | 0.580 | 0.098 | 0.641 | 5.934 | **<0.001** |
| *B. flumineum* (24h) | 0.532 | 0.066 | 0.779 | 8.018 | **<0.001** |
| *B. flumineum* (end) | 0.004 | 0.016 | 0.032 | 0.247 | 0.805 |
| Phytoplankton |  |  |  |  |  |
| F<sub>0</sub> (week 1) | 1.000 |  | 0.361 |  |  |
| F<sub>0</sub> (week 2) | 1.583 | 0.609 | 0.791 | 2.601 | **0.009** |
| F<sub>0</sub> (week 4) | 1.836 | 0.859 | 0.436 | 2.137 | **0.033** |
| F<sub>0</sub> (week 8) | 1.634 | 0.954 | 0.296 | 1.713 | 0.087 |
| QY (week 1) | 2.140 | 0.977 | 0.243 | 2.191 | **0.028** |
| QY (week 2) | 6.357 | 2.443 | 0.793 | 2.602 | **0.009** |
| QY (week 4) | 3.942 | 1.832 | 0.443 | 2.152 | **0.031** |
| QY (week 8) | 4.602 | 2.266 | 0.394 | 2.031 | **0.042** |
| Periphyton F<sub>0</sub> |  |  |  |  |  |
| F<sub>0</sub> (week 1) | 1.000 |  | 0.560 |  |  |
| F<sub>0</sub> (week 2) | 0.966 | 0.296 | 0.536 | 3.262 | **0.001** |
| F<sub>0</sub> (week 4) | 0.900 | 0.345 | 0.403 | 2.612 | **0.009** |
| Periphyton QY |  |  |  |  |  |
| QY (week 1) | 1.000 |  | 0.300 |  |  |
| QY (week 2) | 3.555 | 2.064 | 0.431 | 1.722 | 0.085 |
| QY (week 4) | 2.235 | 1.233 | 0.506 | 1.813 | 0.070 |
| Snail density |  |  |  |  |  |
| *Bi. glabrata* eggs | 1.000 |  | 0.722 |  |  |
| *Bi. glabrata* hatch. | 6.243 | 1.373 | 0.583 | 4.548 | **<0.001** |
| *Bi. glabrata* adults | 1.756 | 0.287 | 0.776 | 6.120 | **<0.001** |
| *Bu. truncatus* eggs | 0.478 | 0.131 | 0.391 | 3.476 | **0.001** |
| *Bu. truncatus* hatch. | 1.536 | 0.199 | 0.963 | 7.703 | **<0.001** |
| *Bu. truncatus* adults | 2.732 | 0.353 | 0.966 | 7.732 | **<0.001** |
| *H. cubensis* eggs | 0.574 | 0.118 | 0.487 | 4.884 | **<0.001** |
| *H. cubensis* hatch. | 1.431 | 0.215 | 0.842 | 6.671 | **<0.001** |
| *H. cubensis* adults | 2.808 | 0.384 | 0.918 | 7.309 | **<0.001** |
| Composite variables | Estimate | Std.err | Standardized estimate | Z-value | P (>\|z\|) |
| Algal production |  |  |  |  |  |
| Phytoplankton | 1.000 |  | 1.149 |  |  |

| Periphyton F0 | -0.044 | 0.138 | -0.111 | -0.321 | 0.749 |
| --- | --- | --- | --- | --- | --- |
| Periphyton QY | 1.705 | 0.635 | 0.999 | 2.685 | **0.007** |
| Covariances | Estimate | Std.err | Standardized estimate | Z-value | P (>\|z\|) |
| Predator mortality |  |  |  |  |  |
| *B. flumineum* (24h) ~<br>*B. flumineum* (end) | 0.005 | 0.002 | 0.335 | 2.507 | **0.012** |
| Phytoplankton |  |  |  |  |  |
| F0(wk 1) ~ QY(wk 1) | 0.328 | 0.085 | 0.589 | 3.841 | **<0.001** |
| F0(wk 4) ~ QY(wk 4) | 0.544 | 0.128 | 0.712 | 4.263 | **<0.001** |
| F0(wk 8) ~ QY(wk 8) | 0.819 | 0.219 | 0.573 | 3.733 | **<0.001** |
| Snail density |  |  |  |  |  |
| *Bu. truncatus* eggs ~<br>*H. cubensis* eggs | 0.006 | 0.001 | 0.637 | 4.145 | **0.000** |
| *Bi. glabrata* eggs ~<br>*Bu. truncatus* eggs | 0.004 | 0.001 | 0.478 | 3.309 | **0.001** |
| *Bi. glabrata* eggs ~<br>*H. cubensis* eggs | 0.003 | 0.001 | 0.343 | 2.490 | **0.013** |
| *Bu. truncatus* hatch ~<br>*H. cubensis* end | -0.002 | 0.001 | -0.558 | -3.348 | **0.001** |
| *Bi. glabrata* hatch ~<br>*H. cubensis* hatch | 0.025 | 0.009 | 0.408 | 2.867 | **0.004** |
| Periphyton F0 ~<br>Periphyton QY | -0.001 | 0.000 | -0.396 | -2.853 | **0.004** |
| Regressions | Estimate | Std.err | Standardized estimate | Z-value | P(>\|z\|) |
| Predator mortality~ |  |  |  |  |  |
| Chlorpyrifos | 1.050 | 0.008 | 0.998 | 133.140 | **<0.001** |
| Phytoplankton~ |  |  |  |  |  |
| Atrazine | -0.059 | 0.022 | -0.201 | -2.735 | **0.006** |
| Fertilizer | 0.234 | 0.022 | 0.793 | 10.806 | **<0.001** |
| Phytoplankton F0~ |  |  |  |  |  |
| Atrazine | -0.274 | 0.027 | -0.424 | -10.303 | **<0.001** |
| Fertilizer | 0.330 | 0.025 | 0.510 | 13.104 | **<0.001** |
| At*Fe | 0.571 | 0.037 | 0.692 | 15.597 | **<0.001** |
| Periphyton QY~ |  |  |  |  |  |
| Atrazine | 0.123 | 0.006 | 0.816 | 19.751 | **<0.001** |
| Fertilizer | -0.069 | 0.006 | -0.458 | -11.082 | **<0.001** |
| Snail density~ |  |  |  |  |  |
| Predator mortality | 0.164 | 0.021 | 0.977 | 7.838 | **<0.001** |
| Algal production | 0.081 | 0.029 | 0.118 | 2.780 | **<0.001** |

**Extended Data Table 2.** Results of a zero-inflated Poisson model on density of infected *Biomphalaria glabrata* at the end of the experiment. The zero-inflated portion of the model included only the intercept and crayfish survival as no other predictor variables were significant and model selection indicated the simpler model was a better fit (ΔAICc >2). Analysis was performed only for infected *Bi. glabrata* as too few infected *Bu. truncatus* were present at the end of the experiment. At = atrazine; Ch = chlorpyrifos; Fe = fertilizer.

| Term | Coefficient | Std. Error | z-value | P |
| --- | --- | --- | --- | --- |
| intercept | -0.23 | 1.14 | -0.20 | 0.842 |
| *Bi. glabrata* density | 0.007 | 0.002 | 4.50 | **<0.001** |
| *P. alleni* survival | 3.03 | 3.84 | 0.79 | 0.429 |
| Block2 | 2.17 | 1.12 | 1.93 | 0.054 |
| Block3 | -3.58 | 7.61 | -0.47 | 0.638 |
| Block4 | 2.47 | 1.12 | 2.20 | **0.028** |
| Block5 | 0.36 | 1.43 | 0.25 | 0.802 |
| At | -2.38 | 1.70 | -1.40 | 0.162 |
| Ch | -1.74 | 1.51 | -1.16 | 0.247 |
| Fe | -17.09 | 145.05 | -0.12 | 0.906 |
| At:Ch | 2.43 | 1.79 | 1.36 | 0.175 |
| At:Fe | 16.89 | 144.90 | 0.12 | 0.907 |
| Ch:Fe | 16.16 | 145.05 | 0.11 | 0.911 |
| At:Ch:Fe | -16.38 | 144.90 | -0.11 | 0.910 |

Zero-inflation model coefficients (binomial with logit link)
| Term | Coefficient | Std. Error | z-value | P |
| --- | --- | --- | --- | --- |
| Intercept | -1.66 | 0.83 | -2.00 | **0.046** |
| *P. alleni* survival | 1.78 | 0.58 | 3.09 | **0.002** |

**Extended Data Table 3.** Results of generalized linear mixed model on infection prevalence of *Schistosoma mansoni* in *Biomphalaria glabrata*. Number of infected *Bi. glabrata* versus number of uninfected *Bi. glabrata* in each replicate was modeled using a beta binomial distribution and fixed effects of all predictor variables. At = atrazine; Ch = chlorpyrifos; Fe = fertilizer.

| Term | Coefficient | Std. Error | z-value | P |
| --- | --- | --- | --- | --- |
| intercept | -3.85 | 1.27 | -3.03 | **0.002** |
| At | 0.36 | 1.29 | 0.28 | 0.780 |
| Ch | -0.32 | 1.14 | -0.28 | 0.777 |
| Fe | -12.48 | 1712.60 | -0.01 | 0.994 |
| At:Ch | -0.28 | 1.47 | -0.19 | 0.847 |
| At:Fe | 12.75 | 1712.60 | 0.01 | 0.994 |
| Ch:Fe | 11.48 | 1712.60 | 0.01 | 0.995 |
| At:Ch:Fe | -11.94 | 1712.60 | -0.01 | 0.994 |
| Block 2 | 0.69 | 0.86 | 0.80 | 0.424 |
| Block 3 | 0.49 | 1.03 | 0.47 | 0.635 |
| Block 4 | 0.19 | 0.91 | 0.21 | 0.831 |
| Block 5 | 0.20 | 0.90 | 0.23 | 0.822 |

**Extended Data Table 4.** Results of generalized linear mixed model on *S. mansoni* cercaria shedding rates from infected *Bi. glabrata*. The number of cercariae shed per hour was modeled using a negative binomial distribution. At = atrazine; Ch = chlorpyrifos; Fe = fertilizer.

| Main effects of agrochemicals |  |  |  |  |
| --- | --- | --- | --- | --- |
| Term | Coefficient | Std. Error | z-value | P |
| Intercept | -0.498 | 1.549 | -0.32 | 0.750 |
| Days post-exposure | 0.138 | 0.035 | 4.01 | **<0.001** |
| At | -0.175 | 0.275 | -0.64 | 0.520 |
| Ch | -0.014 | 0.315 | -0.04 | 0.970 |
| Fe | -0.106 | 0.274 | -0.39 | 0.700 |

| Interaction between atrazine and fertilizer |  |  |  |  |
| --- | --- | --- | --- | --- |
| Term | Coefficient | Std. Error | z-value | P |
| Intercept | -1.262 | 1.959 | -0.64 | 0.519 |
| Days post-exposure | 0.161 | 0.047 | 3.45 | **<0.001** |
| At | -0.341 | 0.370 | -0.92 | 0.356 |
| Fe | -0.321 | 0.394 | -0.81 | 0.415 |
| At:Fe | 0.409 | 0.564 | 0.73 | 0.468 |

**Extended Data Table 5.** Mean percent of *Schistosoma mansoni* cercariae that were dead (standard error of the mean in parentheses) after 2, 4, 8, 12, or 24 h of exposure to solvent control or the estimated environmental concentration of atrazine, chlorpyrifos, or fertilizer. (see Methods for actual concentrations; *n* = 6). Significant treatment effects are in bold. There were no significant effects of treatment on cercarial survival when including time since agrochemical exposure as a predictor variable. When analyzing data from each time point independently, there were no significant effects of treatment at or before 12 h. Given that *S. mansoni* are only infective for approximately 12 h^49^, survival differences after this time period are less ecologically relevant than before it.

| Treatment | 2 h | 4 h | 8 h | 12 h | 24 h | Mean |
| --- | --- | --- | --- | --- | --- | --- |
| Atrazine | 2.78 (2.78) | 8.33 (8.33) | 4.83 (3.88) | 3.27 (2.58) | 74.07 (10.21) | 19.13 (5.96) |
| Chlorpyrifos | 5.56 (5.56) | 1.39 (1.39) | 4.78 (3.36) | 1.52 (1.52) | **86.30 (5.34)** | 19.91 (6.38) |
| Fertilizer | 0.00 (0.00) | 1.19 (1.19) | 6.22 (4.06) | 9.52 (7.06) | **85.57 (7.83)** | 20.99 (6.62) |
| Solvent | 2.38 (2.38) | 1.85 (1.85) | 1.39 (1.39) | 12.00 (4.66) | 55.56 (15.91) | 14.63 (4.98) |

**Extended Data Table 6.** Results of generalized linear mixed model on *S. mansoni* and *S. haematobium* egg viability. The number of hatched eggs was modeled using a beta binomial distribution. At = atrazine; Ch = chlorpyrifos; Fe = fertilizer.

| *Schistosoma mansoni* |  |  |  |  |
| --- | --- | --- | --- | --- |
| Term | Coefficient | Std. Error | z-value | P |
| Intercept | -2.301 | 0.230 | -10.02 | <0.001 |
| At | -0.109 | 0.272 | -0.40 | 0.689 |
| Ch | -0.294 | 0.289 | -1.02 | 0.310 |
| Fe | 0.009 | 0.262 | 0.03 | 0.973 |
| At:Ch | 0.506 | 0.418 | 1.21 | 0.226 |
| At:Fe | 0.358 | 0.404 | 0.88 | 0.376 |
| Ch:Fe | 0.037 | 0.453 | 0.08 | 0.936 |
| At:Ch:Fe | -1.333 | 0.701 | -1.90 | 0.057 |

| *Schistosoma haematobium* |  |  |  |  |
| --- | --- | --- | --- | --- |
| Term | Coefficient | Std. Error | z-value | P |
| Intercept | -3.798 | 0.884 | -4.92 | <0.001 |
| At | 0.066 | 0.431 | 0.15 | 0.880 |
| Ch | -0.175 | 0.455 | -0.38 | 0.700 |
| Fe | -0.200 | 0.467 | -0.43 | 0.670 |
| At:Ch | 0.761 | 0.643 | 1.18 | 0.240 |
| At:Fe | -0.701 | 0.790 | -0.89 | 0.370 |
| Ch:Fe | 1.159 | 0.920 | -1.26 | 0.210 |
| At:Ch:Fe | 1.144 | 1.255 | 0.91 | 0.360 |

**Extended Data Table 7.**
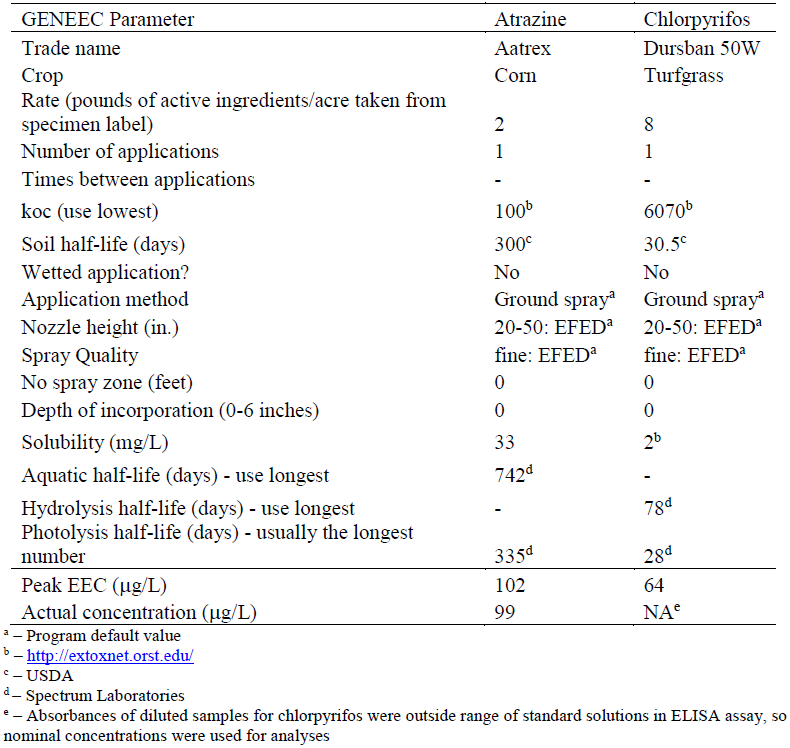
Parameters used for calculation of peak estimated environmental concentrations (EECs).

**Extended Data Table 8.** Results of generalized linear model on *Bi. glabrata* and *Bu. truncatus* densities in response to *Hydrilla verticillata* biomass and the presence or absence of crayfish predators. The number of snails in each mesocosm at the end of the experiment was modeled using a Poisson distribution with interaction terms between the biomass of *Hydrilla* in each tank at the end of the experiment, the presence or absence of predators, their interaction, and spatial block.

| *Biomphalaria glabrata* |  |  |  |  |
| --- | --- | --- | --- | --- |
| Term | Coefficient | Std. Error | z-value | P |
| Intercept | 4.098 | 0.045 | 90.90 | <0.001 |
| *Hydrilla* biomass | 0.007 | <0.001 | 8.60 | <0.001 |
| Crayfish presence | -0.966 | 0.045 | -21.59 | <0.001 |
| *Hydrilla* *crayfish | -0.083 | 0.015 | -5.41 | <0.001 |
| Block2 | 0.297 | 0.053 | 5.57 | <0.001 |
| Block3 | 0.124 | 0.054 | 2.29 | 0.022 |
| Block4 | 1.244 | 0.047 | 26.63 | <0.001 |
| Block5 | -1.532 | 0.086 | -17.71 | <0.001 |

| *Bulinus truncatus* |  |  |  |  |
| --- | --- | --- | --- | --- |
| Term | Coefficient | Std. Error | z-value | P |
| Intercept | 4.614 | 0.034 | 134.80 | <0.001 |
| *Hydrilla* biomass | 0.014 | <0.001 | 19.80 | <0.001 |
| Crayfish presence | -0.642 | 0.043 | -15.08 | <0.001 |
| *Hydrilla* *crayfish | -0.442 | 0.051 | -8.68 | <0.001 |
| Block2 | 0.102 | 0.040 | 2.54 | 0.011 |
| Block3 | -0.805 | 0.048 | -16.83 | <0.001 |
| Block4 | -0.416 | 0.045 | -9.26 | <0.001 |
| Block5 | -2.778 | 0.095 | -29.16 | <0.001 |

**Extended Data Table 9.** Model parameter symbology, definitions, values and sources used as reference literature for parameter values.

| Symbol | Definition | Value | Source |
| --- | --- | --- | --- |
| $f_N$ | Per-capita daily fertility rate of snails including survival to detectability | 0.10 | <sup>64</sup> |
| $\varphi_N$ | Density-dependent snail population parameter, roughly the inverse of snail carrying capacity | $10^3$<br>(~50/m<sup>2</sup>) | <sup>64</sup> |
| $\varphi_{N,q}$ | Scalar of the density dependent snail population parameter caused by fertilizer and/or atrazine stimulation of algal resources | See Extended Data Table 10 | This study |
| $\mu_N$ | Natural per-capita daily mortality rate of snails | 0.017 | <sup>64</sup> |
| $\beta$ | Infection probability from man to snail; interpreted as the per-capita daily probability of snail infection given the number of mated female worms ($M$) | $1.63 \times 10^{-5}$ | Fit to epi data |
| $\sigma$ | Per-capita daily conversion rate of exposed to infected snails | 0.025 | <sup>32</sup> |
| $n$ | Exponent of prey density in Holling III functional response | 2 | <sup>61</sup> |
| $\mu_I$ | Additional per-capita daily mortality of infected snails | 0.083 | <sup>32</sup> |
| $\lambda_{lo}$ | Infection probability from snail to man in low transmission season; interpreted as the per-capita daily probability an adult worm establishes within a human host given the number of shedding snails | $3.67 \times 10^{-6}$ | Fit to epi data |
| $\lambda_{hi}$ | Infection probability from snail to man in high transmission season | $2.45 \times 10^{-4}$ | Fit to epi data |
| $\mu_H$ | Per-capita daily mortality rate of adult worms caused by human mortality (assuming lifespan of 60 years) | $4.57 \times 10^{-5}$ | <sup>5</sup> |
| $H$ | Total human population interacting with water contact site | 300 | <sup>5</sup> |
| $k$ | Clumping parameter of the negative binomial distribution of worms within the human population | 0.08 | Estimated from epi data |
| $\mu_W$ | Natural per-capita daily mortality rate of adult worms (assuming lifespan of 3.3 years) | $8.3 \times 10^{-4}$ | <sup>65</sup> |
| $f_P$ | Per-capita daily fertility rate of predator population including survival to effective snail predation | 0.117 | <sup>66</sup> |
| $\varphi_P$ | Predator carrying capacity | 120 (~0.6/m<sup>2</sup>) | This study |
| $\mu_P$ | Natural per-capita daily predator mortality rate | See Extended Data Table 10 | This study |
| $\mu_{P,q}$ | Additional per-capita daily mortality rate caused by insecticide at concentration, $q$ | See Extended Data Table 10 | <sup>37</sup> |
| $\alpha$ | Per capita attack rate of predators on snails at low densities | 0.003 | <sup>23</sup> |
| $T_h$ | Predation saturation parameter; approximately the inverse of the daily maximum snails consumed per *Procambarus clarkii* predator | 0.067 | <sup>67</sup> |

**Extended Data Table 10.** Model parameter distributions included in the Monte Carlo simulation.

| Symbol | Distribution | Agrochemical Treatment | Distribution parameters |
| --- | --- | --- | --- |
| $\varphi_{N,q}$ | Normal | Fertilizer | Mean = 1.16<br>St. dev = 0.52 |
|  |  | Atrazine | Mean = 1.63<br>St. dev = 0.46 |
|  |  | Atrazine & Fertilizer | Mean = 1.50<br>St. dev = 0.28 |
| $\mu_P$ | Beta | Chlorpyrifos absent | $\alpha = 2.81$<br>$\beta = 70.22$ |
| $\mu_{P,q}$ | | Chlorpyrifos present | $\alpha = 50.39$<br>$\beta = 18.28$ |
| $\beta$ | Weighted by likelihood | NA | 95% CI = $1.63 \times 10^{-6} - 3.11 \times 10^{-5}$ |
| $\lambda_{lo}$ | | NA | 95% CI = $3.67 \times 10^{-7} - 6.97 \times 10^{-6}$ |
| $\lambda_{hi}$ | | NA | 95% CI = $9.51 \times 10^{-4} - 4.66 \times 10^{-4}$ |

## References

1. Steinmann, P., Keiser, J., Bos, R., Tanner, M. & Utzinger, J. Schistosomiasis and water resources development: systematic review, meta-analysis, and estimates of people at risk. Lancet Infect. Dis. 6, 411–425 (2006).

2. May, R. M. Parasites, people and policy: infectious diseases and the Millennium Development Goals. Trends Ecol. Evol. 22, 497–503 (2007).

3. Moran, M. et al. Neglected disease research and development: How much are we really spending? PLoS Med. 6, e1000030 (2009).

4. Savaya Alkalay, A. et al. The prawn *Macrobrachium vollenhovenii* in the Senegal River Basin: towards sustainable restocking of all-male populations for biological control of schistosomiasis. PLoS Negl. Trop. Dis. 8, e3060 (2014).

5. Sokolow, S. H. et al. Reduced transmission of human schistosomiasis after restoration of a native river prawn that preys on the snail intermediate host. Proc. Natl. Acad. Sci. U. S. A. 112, 9650–9655 (2015).

6. Southgate, V. R. Schistosomiasis in the Senegal River Basin: before and after the construction of the dams at Diama, Senegal and Manantali, Mali and future prospects. J. Helminthol. 71, 125–132 (1997).

7. Rohr, J. R. et al. Agrochemicals increase trematode infections in a declining amphibian species. Nature 455, 1235–1239 (2008).

8. Johnson, P. T. J. et al. Aquatic eutrophication promotes pathogenic infection in amphibians. Proc. Natl. Acad. Sci. U. S. A. 104, 15781–6 (2007).

9. Halstead, N. T. et al. Community ecology theory predicts the effects of agrochemical mixtures on aquatic biodiversity and ecosystem properties. Ecol. Lett. 17, 932–941 (2014).

10. Tilman, D. et al. Forecasting agriculturally driven global environmental change. Science. 292, 281–284 (2001).

11. Alexandratos, N. & Bruinsma, J. World agriculture towards 2015/2030: the 2012 revision. ESA Working paper No. 12-03 (2012). doi:10.1016/S0264-8377(03)00047-4

12. United Nations, Department of Economic and Social Affairs & Population Division. World Population Prospects: The 2015 Revision, Key Findings and Advance Tables. Working Paper No. ESA/P/WP.241 (2015). doi:10.1017/CBO9781107415324.004

13. Garibaldi, L. A. et al. Farming approaches for greater biodiversity, livelihoods, and food security. Trends Ecol. Evol. 32, 68–80 (2017).

14. Pretty, J. et al. Resource-conserving agriculture increases yields in developing countries. Environ. Sci. Technol. 40, 1114–1119 (2006).

15. World Health Organization. Schistosomiasis. (2017).

16. World Health Organization. Schistosomiasis and soil-transmitted helminthiases: number of people treated in 2015. Wkly. Epidemiol. Rec. 49/50, 585–600 (2016).

17. Hayes, T. B. et al. Pesticide mixtures, endocrine disruption, and amphibian declines: are we underestimating the impact? Environ. Health Perspect. 114, 40–50 (2006).

18. Relyea, R. A. A cocktail of contaminants: how mixtures of pesticides at low concentrations affect aquatic communities. Oecologia 159, 363–376 (2009).

19. Grube, A., Donaldson, D., Kiely, T. & Wu, L. Pesticides Industry Sales and Usage: 2006 and 2007 Market Estimates. (2011).

20. Miensah, E. D., Fianko, J. R. & Adu-Kumi, S. Assessment of lindane and atrazine residues in maize produced in Ghana using gas chromatography-electron capture detector (GC-ECD) and gas spectrometry (GC-MS). J. Environ. Prot. (Irvine,. Calif). 6, 1105– 1117 (2015).

21. Jepson, P. C. et al. Measuring pesticide ecological and health risks in West African agriculture to establish an enabling environment for sustainable intensification Measuring pesticide ecological and health risks in West African agriculture to establish an enabling environme. Philos. Trans. R. Soc. Lond. B. Biol. Sci. 369, 20130491 (2014).

22. Dabrowski, J. M. Development of pesticide use maps for South Africa. S. Afr. J. Sci. 111, 1–7 (2015).

23. Sokolow, S. H., Lafferty, K. D. & Kuris, A. M. Regulation of laboratory populations of snails (*Biomphalaria* and *Bulinus* spp.) by river prawns, *Macrobrachium* spp. (Decapoda, Palaemonidae): implications for control of schistosomiasis. Acta Trop. 132, 64–74 (2014).

24. Hofkin, B. V & Hofinger, D. M. Predation of Biomphalaria and non-target molluscs by the crayfish Procambarus clarkii: implications for the biological control of schistosomiasis. Ann. Trop. Med. Parasitol. 86, 663–670 (1992).

25. Khalil, M. T. & Sleem, S. H. Can the freshwater crayfish eradicate schistosomiasis in Egypt and Africa? J. Am. Sci. 7, 457–462 (2011).

26. Roberts, J. & Kuris, A. M. Predation and control of laboratory populations of the snail Biomphalaria glabrata by the freshwater prawn Macrobrachium rosenbergii. Ann. Trop. Med. Parasitol. 84, 401–412 (1990)

27. Swartz, S. J., De Leo, G. A., Wood, C. L. & Sokolow, S. H. Infection with schistosome parasites in snails leads to increased predation by prawns: implications for human schistosomiasis control. J. Exp. Biol. 218, 3962–3967 (2015).

28. Mkoji, G. M. et al. Impact of the crayfish *Procambarus clarkii* on *Schistosoma haematobium* transmission in Kenya. Am. J. Trop. Med. Hyg. 61, 751–759 (1999).

29. Evers, B. N., Madsen, H., Alle, J. & Road, B. The schistosome intermediate host, *Bulinus nyassanus*, is a &#x2019;preferred&#x2019; food for the cichlid fish, *Trematocranus placodon*, at Cape Maclear, Lake Malawi. Ann. Trop. Med. Parasitol. 100, 75–85 (2006).

30. Madsen, H. & Stauffer, J. R. Density of Trematocranus placodon (Pisces: Cichlidae): A predictor of density of the schistosome intermediate host, Bulinus nyassanus (Gastropoda: Planorbidae), in Lake Malawi. Ecohealth 8, 177–189 (2011).

31. Rohr, J. R. et al. Predator diversity, intraguild predation, and indirect effects drive parasite transmission. Proc. Natl. Acad. Sci. U. S. A. 112, 3008–3013 (2015).

32. Anderson, R. M. & May, R. M. Infectious Diseases of Humans. (Oxford University Press, 1991).

33. Macdonald, G. The dynamics of helminth infections, with special reference to schistosomes. Trans. R. Soc. Trop. Med. Hyg. 59, 489–506 (1965).

34. Lawal-Are, A. O. Comparative biology of the prawns Macrobrachium macrobrachion (Herklots) and Macrobrachium vollenhovenii (Herklots) from two interconnecting fresh/brackish water lagoons in south-west Nigeria. J. Mar. Sci. Res. Dev. 2, 108 (2012).

35. Hofkin, B. V, Koech, D. K., Oumaj, J. & Loker, E. S. The North American crayfish Procambarus clarkii and the biological control of schistosome-transmitting snails in Kenya: Laboratory and field investigations. Biol. Control 1, 183–187 (1991).

36. Ippolito, A. et al. Modeling global distribution of agricultural insecticides in surface waters. Environ. Pollut. 198, 54–60 (2015).

37. Halstead, N. T., Civitello, D. J. & Rohr, J. R. Comparative toxicities of organophosphate and pyrethroid insecticides to aquatic macroarthropods. Chemosphere 135, 265–271 (2015).

38. Rohr, J. R., Halstead, N. T. & Raffel, T. R. The herbicide atrazine, algae, and snail populations. Environ. Toxicol. Chem. 31, 973–974 (2012).

39. Chase, J. M. Strong and weak trophic cascades along a productivity gradient. Oikos 1, 187–195 (2003).

40. Cohen, L. M., Neimark, H. & Eveland, L. K. Schistosoma mansoni: response of cercariae to a thermal gradient. J. Parasitol. 66, 362–364 (1980).

41. Rosseel, Y. lavaan: An R Package for Structural Equation Modeling. J. Stat. Softw. 48, 1– 36 (2012).

42. R Core Team. R: A language and environment for statistical computing. (2015). at&#x003C;http://www.r-project.org/&#x003E;

43. Jackman, S. pscl: classes and methods for R developed in the Political Science Computational Laboratory, Stanford University. (2015).

44. Zeileis, A., Kleiber, C. & Jackman, S. Regression models for count data in R. J. Stat. Softw. 27, 1–25 (2008).

45. Skaug, H. J., Fournier, D., Bolker, B. M., Magnusson, A. & Nielsen, A. Generalized linear mixed models using &#x2019;AD Model Builder&#x2019;. (2016).

46. Fournier, D. A. et al. AD Model Builder: using automatic differentiation for statistical inference of highly parameterized complex nonlinear models. Optim. Methods Softw. 27, 233–249 (2012).

47. Baer, K. N., Ziegenfuss, M. C., Banks, S. D. & Ling, Z. Suitability of high-hardness COMBO meidum for ecotoxicity testing using algae, daphnids, and fish. Bull. Environ. Contam. Toxicol. 63, 289–296 (1999).

48. McMahon, T. A. & Rohr, J. R. Trypan blue dye is an effective and inexpensive way to determine the viability of *Batrachochytrium dendrobatidis* zoospores. Ecohealth 11, 164– 167 (2014).

49. Olivier, L. J. Infectivity of *Schistosoma mansoni* cercariae. Am. J. Trop. Med. Hyg. 15, 882–885 (1966).

50. Dorn, N. J. & Wojdak, J. M. The role of omnivorous crayfish in littoral communities. Oecologia 140, 150–9 (2004).

51. Mar&çal, A. et al. Predator–prey interactions of *Procambarus clarkii* with aquatic macroinvertebrates in single and multiple prey systems. Acta Oecologica 28, 337–343 (2005).

52. Bronmark, C. Interactions between epiphytes, macrophytes and freshwater snails: a review. J. Molluscan Stud. 55, 299–311 (1989).

53. Turner, A. M., Bernot, R. J. & Boes, C. M. Chemical cues modify species interactions: the ecological consequences of predator avoidance by freshwater snails. Oikos 88, 148–158 (2000).

54. Turner, A. M., Turner, S. E. & Lappi, H. M. Learning, memory and predator avoidance by freshwater snails: effects of experience on predator recognition and defensive strategy. Anim. Behav. 72, 1443–1450 (2006).

55. Turner, A. M. Non-lethal effects of predators on prey growth rates depend on prey density and nutrient additions. Oikos 104, 561–569 (2004).

56. Hoverman, J. T., Auld, J. R. & Relyea, R. A. Putting prey back together again: integrating predator-induced behavior, morphology, and life history. Oecologia 144, 481–91 (2005).

57. May, R. M. Togetherness among schistosomes: its effects on the dynamics of the infection. Math. Biosci. 35, 301–343 (1977).

58. Holling, C. S. The components of predation as revealed by a study of small-mammal predation of the European pine sawfly. Can. Entomol. 91, 293–320 (1959).

59. Diekmann, O., Heesterbeek, J. A. P., Roberts, M. G., Heesterbeek, H. & Roberts, M. G. The construction of next-generation matrices for compartmental epidemic models. J. R. Soc. Interface 7, 873–885 (2010).

60. Hassell, M. P., Lawton, J. H. & Beddington, J. R. Sigmoid functional responses by invertebrate predators and parasitoids. J. Anim. Ecol. 46, 249–262 (1977).

61. Real, L. A. The kinetics of functional response. Am. Nat. 111, 289–300 (1977).

62. Denny, M. Buzz Holling and the functional response. Bull. Ecol. Soc. Am. 95, 200–203 (2014).

63. Baxter, L. R. et al. Atrazine does not affect algal biomass or snail populations in microcosm communities at environmentally relevant concentrations. Environ. Toxicol. Chem. 30, 1689–1696 (2011).

64. Woolhouse, M. E. J. & Chandiwana, S. K. Population biology of the freshwater snail *Bulinus globosus* in the Zimbabwe highveld. J. Appl. Ecol. 27, 41–59 (1990).

65. Goddard, M. J. & Jordan, P. On the longevity of *Schistosoma mansoni* in man on St. Lucia, West Indies. Trans. R. Soc. Trop. Med. Hyg. 74, 185–191 (1980).

66. Cervantes-Santiago, E., Hern&ández-Vergara, M. P., P&érez-Rostro, C. I. & Olvera-Novoa, M. A. Reproductive performance of the crayfish *Procambarus (Austrocambarus) acanthophorus* Villalobos 1948 under controlled conditions. Aquaculture 308, 66–70 (2010).

67. Hofkin, B. V, Mkoji, G. M., Koech, D. K. & Loker, E. S. Control of schistosome-transmitting snails in Kenya by the North American crayfish Procambarus clarkii. Am. J. Trop. Med. Hyg. 45, 339–44 (1991).

